# Marine-freshwater prokaryotic transitions require extensive changes in the proteome

**DOI:** 10.1101/544619

**Authors:** Pedro J. Cabello-Yeves, Francisco Rodriguez-Valera

## Abstract

The comparison of microbial genomes found in either freshwater or marine habitats indicated that in some cases (*Synechococcus* and *Ca*. Pelagibacter) there were notable differences in the global isoelectric point (pI) of proteins. We have analysed global metagenomic proteomes and have added more prokaryotes to extend the pI comparison. Without exception, in a set that included archaea and multiple bacterial phyla, the proteome pI distribution varied, with more acidic values in marine and neutral/basic in freshwater microbes. Four pairs of highly related prokaryotes of marine and freshwater origin revealed marked differences manifested mostly in the residues located at the protein surface. This study has also shown that the magnitude of the change depended on protein location (secreted > cytoplasmic > transmembrane) and affected proteins encoded at both core and flexible genome. Our results point to a very extensive variation taking place in microbes when they move from marine (salt-rich) to freshwater habitats. These adaptations would require long evolutionary times to produce changes involving many genes in the core genome. They also point to significant differences in the physiology, probably at the level of membrane functioning, bioenergetics, intracellular ion concentration and pH (or all of them).

## Introduction

One classic conundrum of microbiology, or actually of biology at large, is the marked borderline that exists between freshwater and marine habitats [1]. Although aquatic environments share many features and ecological parameters, the microbes found throughout both systems have different characteristics at several levels. First, although the major microbial taxa have representation in both, the proportions of each are very different, Actinobacteria being one notorious example of a phylum that is more abundant in freshwater systems [2, 3] or Alphaproteobacteria class having higher representation in marine water bodies, particularly SAR11 clades I/II (*Ca*. Pelagibacter) [4]. Second, although it might be an artefact of lack of coverage, there are specific taxa that appear to be altogether absent in one of the groups of habitats regardless of how abundant they are in the other. Some relevant examples are acI Actinobacteria [5] and LD12 Alphaproteobacteria [6–8], which dominate freshwater habitats, or *Prochlorococcus* species found only in marine ecosystems [9]. The explanation for such differences is not clear considering that the pelagic habitats of off-shore marine waters and oligotrophic lakes are overall similar. There are two obvious differences, salinity and the influence of terrestrially derived organic matter in lakes, that does not reach off-shore marine waters [1]. There are reports of multiple marine clades being detected albeit in small numbers in freshwater habitats [10–12] and the opposite is true for marine regions neighboring the continents, particularly near large estuaries like the Amazon on the Atlantic coast of Brazil [13]. Thus, the differences cannot be explained by physical isolation between these aquatic environments.

One problem to understand the real differences between these two major types of aquatic systems is the enormous diversity within habitats, particularly freshwater lakes, in their trophic status (from oligotrophic to highly eutrophic), nutrient concentrations (phosphorous limitation) and other environmental parameters, all of them having profound implications in the taxonomic composition. Recently, we were involved in the first metagenomic study of Lake Baikal, Siberia, Russia [14]. This is the largest and deepest (max. 1600 m, average 1300 m) lake on Earth [15], ultraoligotrophic and with relatively little influence from terrestrial sources (all features that make it similar to marine off-shore waters) while having very low salt content (dominated by Ca^2+^ and HCO_3_^−^, being particularly low in Na^+^ and K^+^) [16–18]. Interestingly, we found some groups with close relatives among bona fide marine lineages, including the first freshwater *Pelagibacter*-like metagenome assembled genome (MAG) inside the typically marine clades I/II [14]. In previous studies, we compared the pI patterns of SAR11 [14] and *Synechococcus* [19] with their marine close relatives and we noticed significant differences in the acidity and basicity of their proteomes, which were consistently found between the marine and their freshwater counterparts.

The variations in the global pI plots in the proteomes of bacteria or archaea depend on the amino acid overall charge, that has important implications on protein structure and properties [20]. It is generally accepted that prokaryotic genomes have a bimodal shape with two maxima [21], one at acidic pH corresponding largely to dissolved proteins (cytoplasmic or secreted) and one at basic pH of the membrane proteins that have a basic (positively charged) domain intracellularly to facilitate the generation of a proton motive force. In between these two peaks there is a minimum at ca. neutral values that corresponds to the intracellular pH at which proteins of equivalent pI value would be the least soluble. In salt-in halophiles the alkaline peak nearly disappears because they have a large amount of intracellular K^+^ [22]. The adaptation to hypersaline environments (containing much more than the salinity of seawater) leads to these changes in their inhabitants (halophiles) and has been known for long [23]. Thus, hyperhalophiles such as *Haloquadratum walsbyi* or *Salinibacter ruber* have their proteomes severely displaced to acidic values [22]. However, marine bacteria and archaea are expected to be salt-out strategists i.e. that keep most inorganic salts (particularly Na^+^) outside the cell, maintaining a relatively salt free cytoplasm [22].

The large change in these values detected in the freshwater microbes mentioned above made us wonder if these could be a general phenomenon and what could be the underlying reason for such broad deviation. Although there is a current database with pI calculations and amino acid properties for more than >5000 bacteria and archaea [24], there are no studies comparing bona fide freshwater and salt-adapted microbial proteomes. Here, filling this gap, we have analyzed in detail some specific cases when closely related microbes by whole genome comparisons have been retrieved from marine and freshwater habitats; furthermore, there is metagenomic evidence showing that they are actually adapted to live in either one or the other environment. We have also dissected these pI values depending on the localization of the proteins and used models available to determine whether there was a preferential location of the charges. Our data confirm that indeed the proteins regardless of location in the cell accumulate more negative charges in those prokaryotes coming from marine environments, corresponding to a significant deviation in the amino acidic composition that among other consequences imply a large sequence variation and requires long evolutionary times. There is no known example of a single species (microbes with >95% average nucleotide identity, ANI) being found at both aquatic systems.

## Results

### Global pIs of metaproteomes from different aquatic habitats

A global metagenomic approach was first used to assess if the changes in pI could be detected in the microbial community as a whole. Specifically, we used metagenomic datasets from freshwater (Tous reservoir and Lake Baikal), brackish (Caspian Sea) and marine (Mediterranean Sea) systems from similar depths, which were assembled and annotated, taking all proteins from contigs > 5 Kb for the analysis and obtaining sets of more than 85,000 proteins (Fig. 1). Interestingly, three peaks were observed located at nearly identical pH values (4.5, 6.8 and 9.8) for the different aquatic habitats in spite of the large differences in salinity (ca. 0.05, 2 and 3.8 % respectively) and community structure [14, 25, 26]. A major difference was observed in the acidic peak, with brackish and marine environments having higher relative frequencies of lower pIs, compared to their freshwater counterparts. On the other hand, it is observable a higher peak of neutral pIs in freshwater systems. Finally, the distribution of basic pIs (8-9.5) is also higher in freshwater systems. The relative frequencies of pIs from 9.5-14 are a little higher in the Mediterranean Sea, but this could just be a result of a taxonomy bias due to the abundance of SAR11 clade microbes in this sample (see below). However, these changes could just reflect the variation in the community structure i.e. very different taxonomic composition depending on the habitat.

**Fig. 1.**
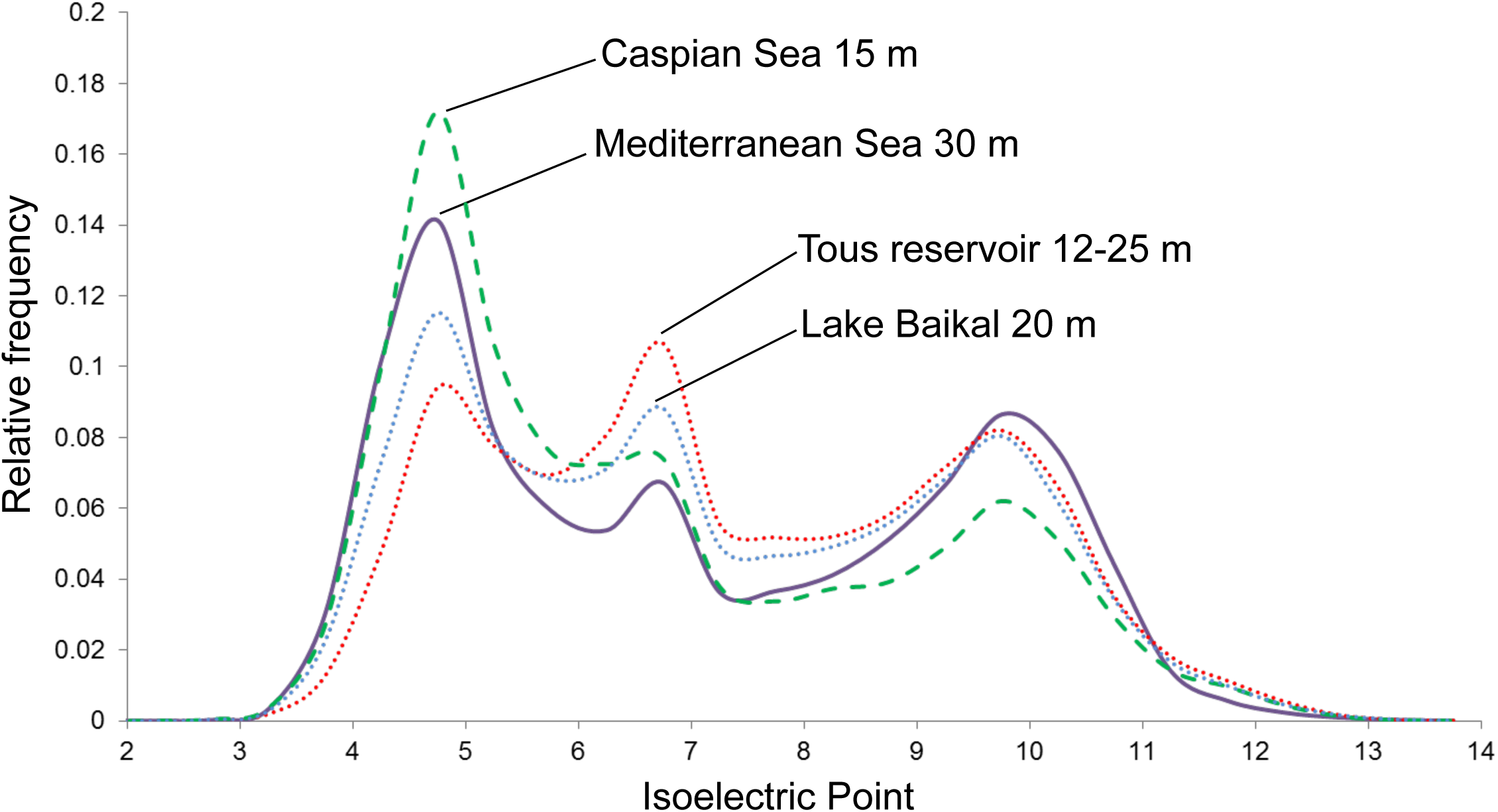
Metaproteome isoelectric point versus relative frequency plot of marine (Mediterranean Sea, 30 m), brackish (Caspian Sea, 15 m) and freshwater (Tous reservoir, 12-25 m and Lake Baikal, 20 m) habitats.

### Overall pI pattern in different phyla

To assess if the differences in the global pI distributions were due to the taxonomic bias we selected a total of 74 prokaryotes from public database (Supplementary Data Set S1) and compared their overall pI values. We used examples of bona fide freshwater, brackish and marine microbes, some of them retrieved from the environments compared in Fig. 1. We took representatives from class Alphaproteobacteria (SAR11, *Roseobacter* and Rhodospirillaceae), order Betaproteobacteriales, Chloroflexi, Planctomycetes, Verrucomicrobia, Cyanobacteria (*Synechococcus*/*Cyanobium*), Actinobacteria, Bacteroidetes and Thaumarchaeota phyla (Fig. 2 and Figs. S1-S6 from Supplementary Material). In all cases there appears to be a significant difference in freshwater, brackish and marine proteomes, as depicted in Figs. 1 and 2, being relatively straightforward the identification of the different microbes according to their origin.

**Fig. 2.**
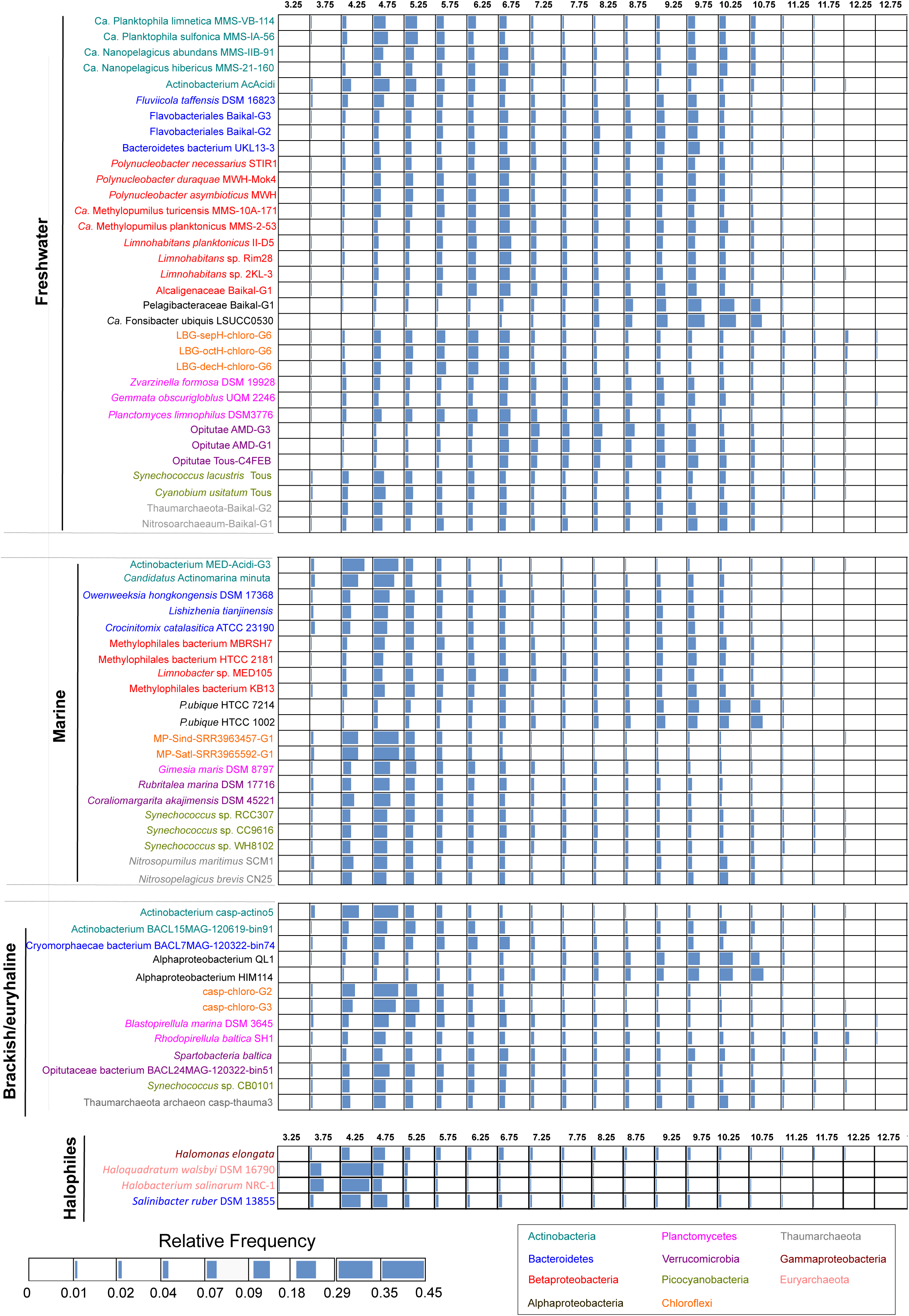
Whole-proteome heatmap-bar isoelectric point versus relative frequency plot of freshwater, marine, brackish and halophilic selected prokaryotes. Genomes are color-coded according to their taxonomic affiliation and divided according to their origin.

First, halophiles present a single acid peak at low pIs (highest among the compared microbes). Second, brackish and marine species tend to show bimodal patterns and display a higher peak of acidic proteins compared to freshwaters, to some extent such as SAR11 (Fig. S1 from Supplementary Material), which always present a higher peak of basic proteins independently of their origin. Third, it is particularly remarkable the high peak of neutral proteins (with pIs ranging from 6 to 8) in some freshwater species, whilst this peak is very low or absent in salt-adapted species. Finally, some freshwater microbes display a multi-modal pI plot (normally consisting of three identifiable peaks), as in the case of Thaumarchaeota, Betaproteobacteriales, Flavobacteriales, Verrucomicrobia or Planctomycetes (Figs. S2, S5 and S6 from Supplementary Material). Our data facilitates an identification of microbes coming from salty ecosystems only by looking at the more acidic pI these microbes display. Hence, it could be stablished as a rule of thumb to infer the preferred habitat of microbes from the same families and genus (and, to some extent, phyla) based on the pI distribution of their proteomes.

### Pairwise comparisons of close phylogenetic neighbors

We selected couples of microbes that are closely associated phylogenetic neighbors but are truly freshwater or marine inhabitants. In these cases, the effect of the taxonomic distance was reduced to the minimum presently available in databases. Thus, we could compare two species from the family Nitrosopumilaceae (*Nitrosoarchaeum* sp. Baikal-G1, MAG, vs *Nitrosopumilus maritimus* SCM1, culture), two SAR11 members from family Pelagibacteraceae (Pelagibacteraceae bacterium Baikal-G1, MAG, vs *Pelagibacter ubique* HTCC7214, culture), two picocyanobacteria from the order Synechococcales (*Synechococcus* sp. RCC307, culture, vs *Synechococcus lacustris* Tous, culture) and finally two species from the family Methylophylaceae (Methylophilales bacterium MBRS-H7, culture vs *Methylopumilus planktonicus* MMS-2-53, culture). We chose microbes with similar cell and genome sizes displaying similar metabolic and ecological roles in the environment to reduce to the minimum other factors. Values of Average Nucleotide Identity (ANI), Average Amino Acid Identity (AAI), 16S rRNA identity and Percentage of conserved Proteins (POCP) were also calculated for each pair. We divided the proteome into three categories: cytoplasmic and inner membrane proteins that are submitted to the cytoplasmic environment, proteins with transmembrane domains, and secreted (with signal peptide), i.e. exposed to the extracellular environment. The average pIs were also calculated for these three categories. The differences between freshwater and marine microbes appear clear at all levels (Figs. 3-6). Remarkably, we found in these couples of microbes whose ANI varied between 66 and 77 that the AAI values were similar (when not lower) than the nucleotide identity (ANI). This is contrary to the expectations in comparisons of phylogenetic relatives that tend to have more similarity at the level of amino acids than nucleotides [27, 28]. This also indicated a major shift in the composition of amino acids of the core genome (shared genes, see below).

**Fig. 3.**
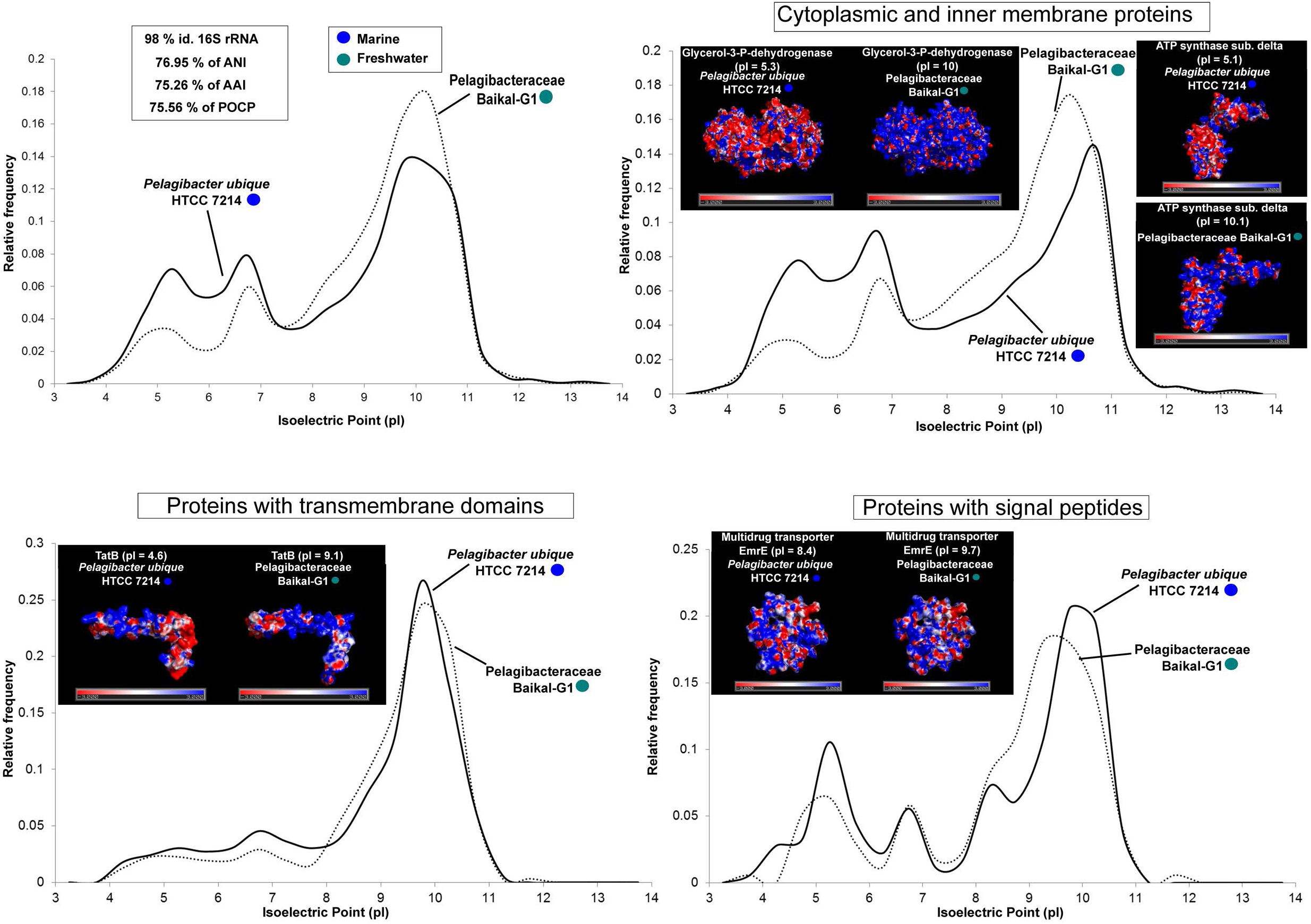
Whole-proteome pI versus relative frequency of *P.ubique* HTCC7214 (marine) and Pelagibacteraceae bacterium Baikal-G1 (freshwater). Tridimensional electrostatic surface potential plots of individual proteins selected for each category. The potentials were mapped from −3 kcal mol^−1^ (red) to +3 kcal mol^−1^ (blue). Values of ANI, AAI, POCP and % of identity of 16S rRNA are also indicated.

**Fig. 4.**
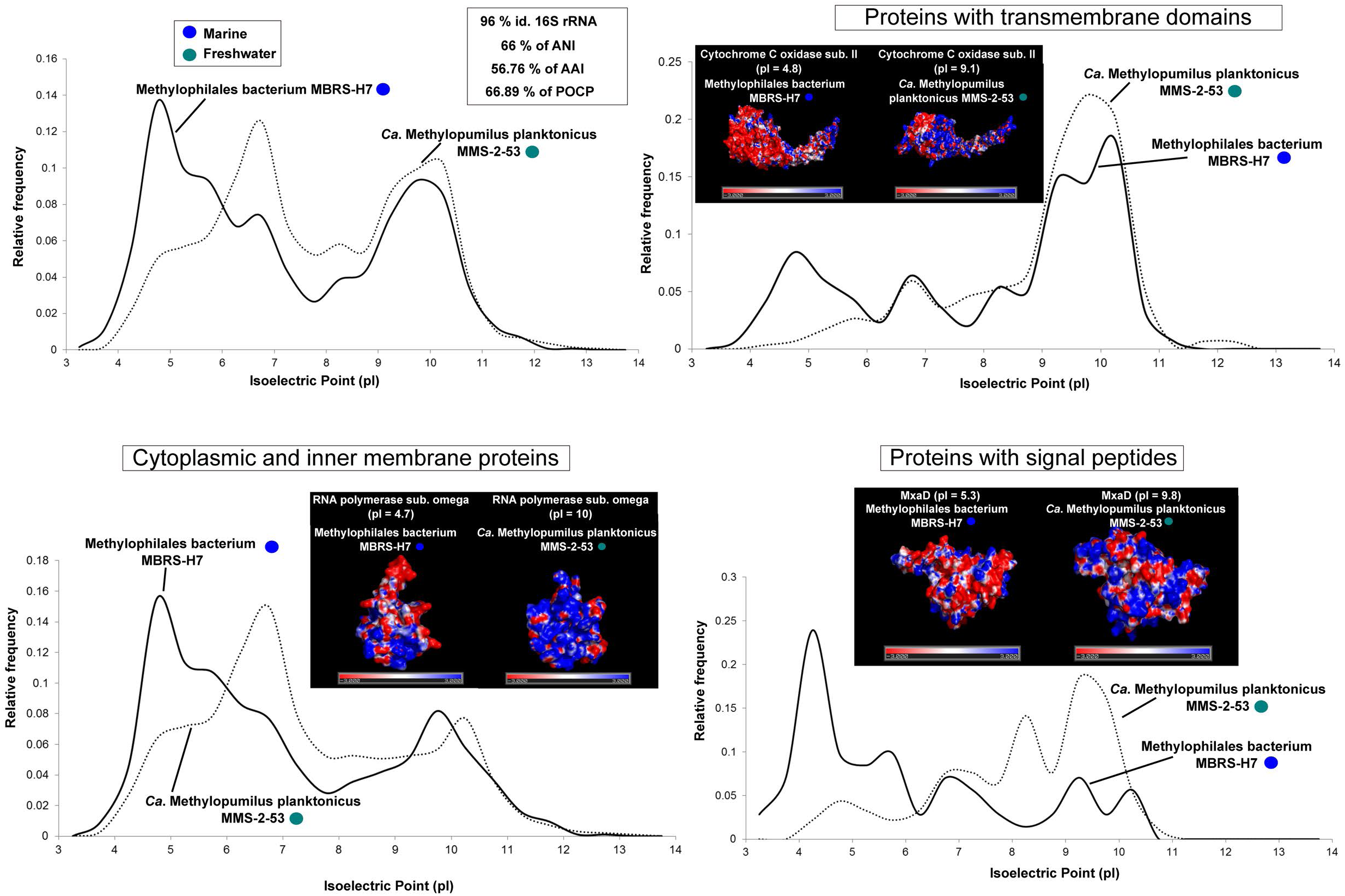
Whole-proteome pI versus relative frequency of *Methylopumilus planktonicus* MMS-2-53 (freshwater) and *Methylophilales bacterium* MBRSH7 (marine). Tridimensional electrostatic surface potential plots of individual proteins selected for each category. The potentials were mapped from −3 kcal mol^−1^ (red) to +3 kcal mol^−1^ (blue). Values of ANI, AAI, POCP and % of identity of 16S rRNA are also indicated.

**Fig. 5.**
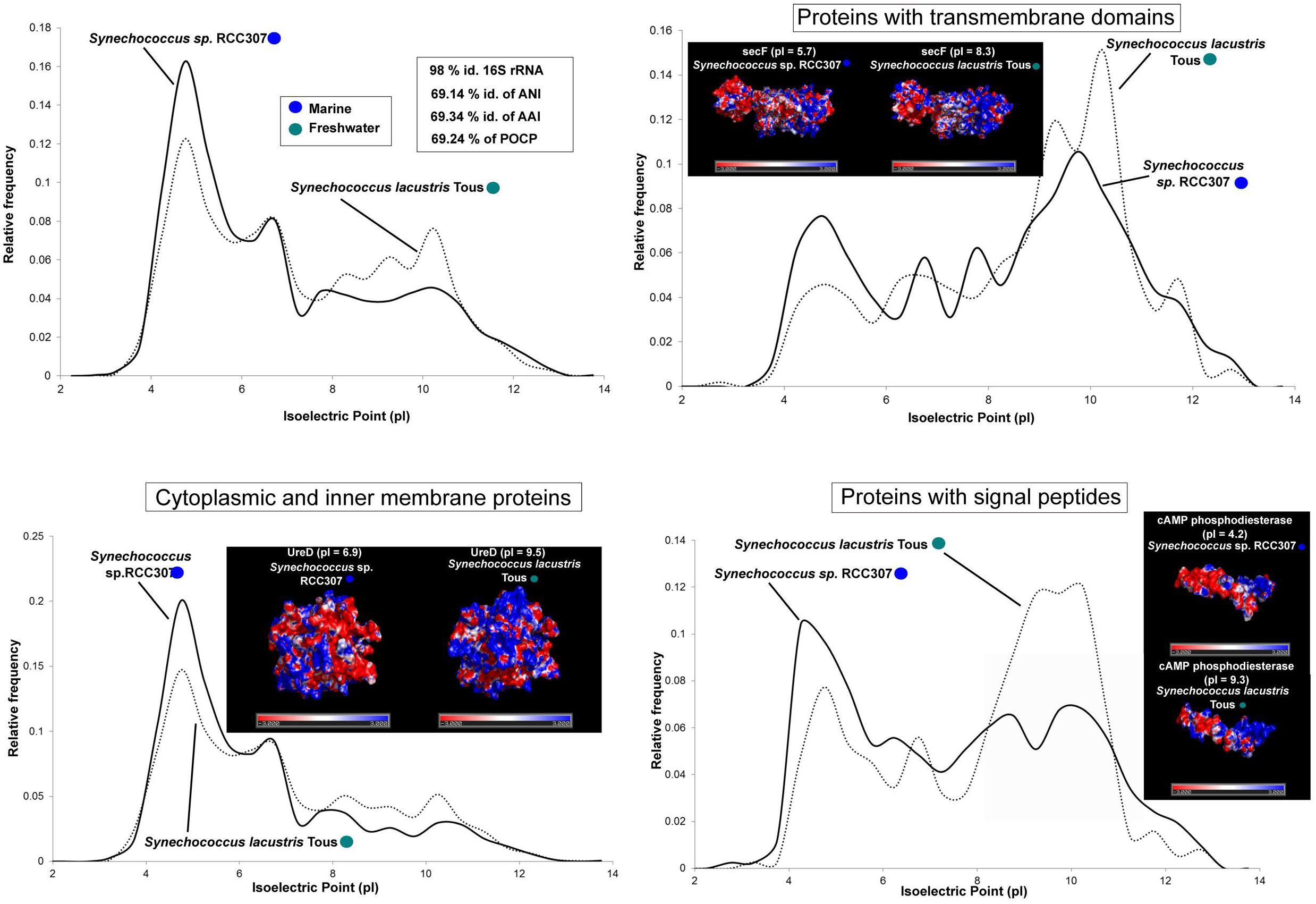
Whole-proteome pI vs relative frequency of *Synechococcus lacustris* Tous (freshwater) and *Synechococcus* sp. RCC307 (marine). Tridimensional electrostatic surface potential plots of individual proteins selected for each category. The potentials were mapped from −3 kcal mol^−1^ (red) to +3 kcal mol^−1^ (blue). Values of ANI, AAI, POCP and % of identity of 16S rRNA are also indicated.

**Fig. 6.**
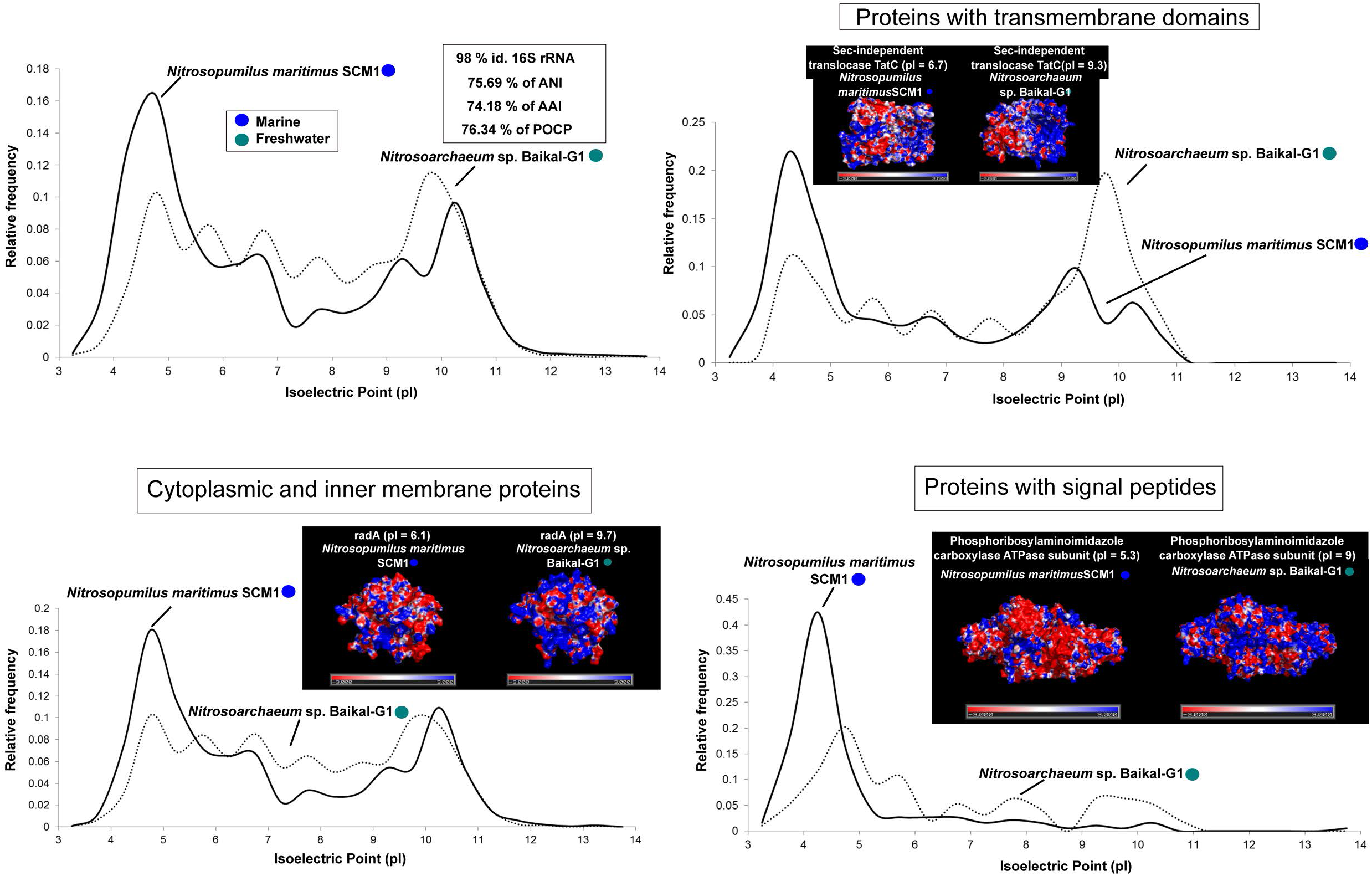
Whole-proteome pI versus relative frequency of *Nitrosopumilus maritimus* SCM1 (marine) and *Nitrosoarchaeum* sp. Baikal-G1 (freshwater). Tridimensional electrostatic surface potential plots of individual proteins selected for each category. The potentials were mapped from −3 kcal mol^−1^ (red) to +3 kcal mol^−1^ (blue). Values of ANI, AAI, POCP and % of identity of 16S rRNA are also indicated.

The pattern of higher acidic proteins is preserved for marine microbes, whilst the neutral to basic predominance is maintained in all freshwater genomes, as observed in the metagenomes or in other less related microbes (see above). We analyzed the case of some individual protein homologs for which there were tridimensional models to assess if the difference in the predicted global pI was concentrated at the level of the protein surface. We chose homologs with a marked difference in the global pI between freshwater and marine representatives; we considered only those proteins with well-stablished tertiary structures (retrieved from SWISS-MODEL). We have observed a higher accumulation of negative charges (acid amino acids) on the surface of proteins from marine microbes, whilst positive electrostatic potentials (with more basic amino acids) are seen in freshwater microbes (Figs. 3-6). Thus, it appears that the electrostatic surface potential between pairs of homologs differs significantly between marine and freshwater species. These differences are even more evident when introducing halophiles into the comparison. Such comparative example within the archaeal domain comprising one halophile (*Haloquadratum walsbyi* DSM 16790), one marine (*Nitrosopumilus maritimus* SCM1) and one freshwater archaeon (*Nitrosoarchaeaum* sp. Baikal-G1) is shown in Fig. S7 from Supplementary Material. These significant variations were apparent in all three categories: cytoplasmic, membrane and secreted proteins. However, the differences in electrostatic surface potential were more marked in the secreted>cytoplasmic>membrane, what could indicate a more radical change in the extracellular than in the intracellular environment between freshwater and marine microbes. We are showing only some individual protein examples for each category/prokaryote, however, all were analyzed and presented substantial differences in their pIs and electrostatic surface potentials (Supplementary Data Set S2).

The variations in the amino acid composition (expressed as Mole% for each microbe) in the compared pairs are noteworthy (Fig. S8 from Supplementary Material). A general trend that is conserved in the four cases is the higher percentage of acid amino acids (aspartic and glutamic) in the marine representatives (from 0.6 to 1.4 % higher). This is in agreement with the overall higher peak of acidic proteome pIs in these salt-adapted microbes. Actually, the global percentage of charged amino acids comprising both acid and basic types (aspartic and glutamic acids, lysine, histidine and arginine), is higher in marine microbes i.e. the acid increase in marine is more accentuated than the basic in freshwater representatives. However, three out of four freshwater microbes analyzed here display higher percentages of basic amino acids compared with their marine relatives. The only exception observed is the case of *S. lacustris* Tous, which presents a nearly identical % of basic amino acids when compared with the marine relative *Synechococcus* RCC307. Nevertheless, there are noticeable differences, for instance, *S.lacustris* shows a higher percentage of lysine (K) residues on its whole proteome, whilst RCC307 compensates this decrease in lysine by having more arginine (R) residues, resulting in practically the same % of total basic amino acids in both genomes. On the other hand, the higher percentage of basic residues in Methylophilales, Pelagibacteraceae and archaeal Nitrosopumilaceae genomes is significant. Nitrosoarchaeum sp. Baikal-G1 presents a higher percentage of all three basic amino acids, compared to its marine relative *Nitrosopumilus maritimus* SCM1. *Ca*. Methylopumilus planktonicus MMS-2-53 displays a higher percentage of arginine and histidine, but slightly lower lysine compared to its marine relative. Hence, a global picture containing these variations in the overall pI plots, electrostatic surface potential of proteins and percentages of basic/acid amino acids among the 4 pairs of microbes could help predicting the freshwater or salt-adapted origin of a novel microbe of unknown source e.g. when analyzing genomes of mixing zones such as estuaries or costal zones in general.

### Pan-genome pI distribution

We also calculated the pIs for each category (core and flexible proteins) to assess if the change in the global proteome might be due to variations affecting homologous proteins as those shown above or it could be an extreme consequence of a change in the differential gene pool present in fresh water or marine microbes i.e. it could be due to the flexible or core genes [29, 30]. Therefore, we analyzed the global pI plots of both components in the four comparisons (Fig. S9 from Supplementary Material). Indeed, marine representatives always had a higher peak of acidic proteins (independently if the coding genes belong to the core or flexible genome component). Actually, there were significant differences in some cases between the pI profiles of the core and flexible genes but not altering significantly those observed in freshwater and marine whole genomes. The flexible genome had in three of the examples a higher basic peak, probably reflecting enrichment in membrane proteins, largely transporters and sensors that are typical components of the flexible genome (involved in habitat-adaptation and niche-occupancy) [31].

## Discussion

Previous studies showed that bacteria and archaea, independently of their origin, presented a bimodal pI pattern, whilst eukaryotes showed a trimodal pattern [21]. Others have shown a multi-modal distribution picture of the pIs in different organisms as a consequence of the chemical properties of the different amino acids (rather than sequence evolution) [20] or as a result of discrete pKr values for the amino acids [32]. Our observations indicate that the pI pattern varies significantly among microbes, having cases of unimodal (halophiles, marine microbes such as Acidimicrobiia, SAR11), bimodal patterns (most of the microbes analyzed here) to multimodal (Thaumachaeaota, Verrucomicrobia, Planctomycetes or Betaproteobacteriales).

The pI of a certain protein is a major indicator of the properties of the macromolecule. It determines the water solubility of the protein as well as the degree of interactions with the chemical environment. Since intracellular pH of most microbes (including alkaliphilic or acidophilic ones) is near neutrality [33, 34] and proteins are less soluble at pH values near their pI, cytoplasmic and secreted proteins, that mostly work in a soluble form, tend to have pIs far from neutrality, mostly acidic [33]. On the other hand, membrane proteins only interact with water in their exposed domains and tend to have alkaline pIs to compensate for the positive charge outside of the membrane created by the proton gradient [35]. The consistent difference that we have detected between freshwater and marine microbes indicates a significant change in one of the two aspects of cell biology, either the intracellular pH or the bioenergetics of the cell (perhaps both). Another factor that likely interacts with the protein charge is the presence of other solutes in the water phase at both sides of the membrane. Most cells maintain a significant concentration of K^+^ cations inside while keeping Na^+^ outside. Typically, marine microbes would need higher intracellular potassium concentrations in order to compensate the sodium ions abundant in the extracellular environment. Depending on the cation concentration soluble proteins need to have more or less negative charges to maintain a proper hydration sphere [36]. This is why halophiles with salt-in strategies must have very acidic soluble proteins [22]. Freshwater must impose limitations to the accessibility to the main cellular cations, particularly Na^+^ that might be limiting in salt-poor environments like Lake Baikal [16]. These conditions could lead to adaptations consisting of less intracellular potassium. It is thus not surprising that a less acidic proteome might be favorable for freshwater microbes. Other predicted physiological differences between the two types of microbes include the preference for H^+^ over Na^+^ based electron or nutrient transport mechanisms [33] but this is unlikely to have effects over the global proteome as described here.

The kind of analysis that we have done in this work has previously been hampered by the lack of close relatives of the same species specialized in living in either freshwater or marine (salts rich) habitats. The study of our metagenome of Lake Baikal [14], the largest freshwater body on Earth that is also similar to marine habitats in most aspects (including extreme oligotrophy and relative little terrestrial influence due to its enormous water mass), revealed the presence of novel and abundant microbes that do not have an allochthonous origin (particularly considering that this lake is very far from the nearest marine water body). However, provided that most of the models that we have used are only available as genomes, it is not feasible to carry out physiological or biophysical experiments that could clarify the meaning of the patterns that we have found. For instance, one crucial point would be determining the intracellular K^+^ concentration, which to date has been only determined in *E.coli* [37], a marine *Pseudomonas* [38] and some halophiles [39]. Similarly, we must first understand how different microbes regulate their cytoplasmic pH in response to environmental changes. However, there is a significant difficulty in measuring the cytoplasmic pH of microbes under growth conditions [34]. Furthermore, some microbes undergo small variations in their cytoplasm’s pH (up to 0.1 units per pH unit change), whilst others such as *E.coli* or *Coxiella burnetti* are subjected to much higher changes [40, 41].

As could be expected, there is a taxonomic factor in the pI patterns, for example, SAR11 clade members tend to have the pI plot displaced towards basic values. Still, even in these cases, the differential value in the freshwater-marine comparison was detectable (i.e. regardless of the pI range always marine have more acidic average values). That streamlined bacteria, independently of their origin, should have a tendency to basicity in their pIs is to be expected (Fig. S10 from Supplementary Material) considering their higher surface/volume ratio, leading to a higher membrane/cytoplasmic proteins ratio. This general pattern was confirmed by the amino acid composition that shows common trends in organisms as phylogenetically distant as *Pelagibacter* and Thaumarchaeota. In marine microbes there was always a decrease in basic amino acids and an increase in acid ones. It was also remarkable that in closely related marine and freshwater microbes AAI was similar and (in most cases) lower than ANI i.e. amino acid similarity is lower than nucleotide identity. This was also observed in freshwater, euryhaline and marine *Synechococcus/Cyanobium* genera [42]. This is the opposite to what is found when comparing similarly distant microbes but living in the same type of aquatic habitat, such as freshwater acI Actinobacteria [5]. The values are consistently AAI<ANI as could be expected from the existence of neutral changes due to the degeneration of the genetic code.

Our work underscores the important changes that a microbe must suffer to get adapted to freshwater from a marine habitat or vice versa. If many (or most) proteins change in their amino acidic composition the amount of changes and evolutionary time involved have to be large and consequently, it is not a simple evolutionary step. Although several studies assured that marine-freshwater transitions tended to be infrequent [43, 44], it has been proven that some close relatives to marine microbes are found in freshwater habitats (SAR11 Pelagibacteraceae, Rhodobacteraceae and Flavobacteria) [10–12, 14]. Therefore, the transition, although demanding, must have happened frequently in the long evolutionary history of microbes. Actually, there are not real geographic or physical barriers separating them since marine microbes are often found (in very small amounts) in terrestrial habitats [45] and fresh water microbes are often transported to the sea e.g. downstream by rivers or lixiviation. The adaptation to be a freshwater or a marine inhabitant is a crucial evolutionary choice that each species makes, likely in its origin as such.

## Conclusions

There is a large change in amino acid composition among microbes depending on whether they live in marine or freshwater habitats. The change can already be detected by relatively low values of AAI (compared to ANI) and is reflected by a major shift in the pI pattern of the cell proteome, with an increase in the acidic peak in the marine microbes and another (albeit more moderate) in the neutral and basic peaks for the freshwater ones. These changes occur also in closely related microbes i.e. they do not reflect a taxonomic bias. Furthermore, we have been able to see changes in individual proteins with 3D models and their overall surface electrostatic potential, indicating that the changes tend to accumulate on the surface of the protein, particularly when they are soluble (cytoplasmic or secreted).

We propose that our results indicate an important change in cell physiology due to the absence of salts in the freshwater habitats. This absence might imply specific requirements of membrane characteristics (membranes could change in composition when exposed to absence of salts in significant amounts since the stability of lipid bilayers could be affected), bioenergetics (differences in the electrochemical gradient across the membrane) intracellular pH (a change in the intracellular pH would modify the solubility of the proteins; if it was higher in freshwater adapted cells it would require the proteins to be more alkaline and less acidic to optimize solubility) or K^+^ concentration (requiring less acidity to compensate the positive charge of intracellular ions), or a combination of these or other components of cell biology which apply throughout the prokaryotic domain, bacteria and archaea.

## Materials and Methods

### Metagenomic datasets and bacterial genomes used in this work

All metagenomic datasets used in this work are publicly available in the NCBI/SRA databases: Mediterranean Sea [26], Caspian Sea [25], Lake Baikal [14], Tous reservoir [19]. All bacterial and archaeal genomes used in this study, together with their accession/Genbank number (NCBI), habitat, isolation/origin, reference, type of genome and phylum are shown in Supplementary Data Set S1. The eight genomes used in the protein-by protein-based comparison were previously published and analyzed by different tools: *Synechococcus lacustris* Tous [42], *Synechococcus* sp. RCC307 [46], *Methylopumilus planktonicus* MMS-2-53 [47], *Methylophilales bacterium* MBRSH7 [48], Pelagibacteraceae bacterium Baikal-G1 [14], *Pelagibacter ubique* HTCC 7214 (ASM70138v1), *Nitrosopumilus maritimus* SCM1 [49] and *Nitrosoarchaeum* sp. Baikal-G1 [14].

### Protein isoelectric point determination

The isoelectric point calculations and amino acid features of each protein and microbe were calculated with the software Pepstats from the EMBOSS package [50]. To determine the pI distribution from metaproteomes we obtained all proteins from the assembled contigs larger than 5 Kb. We used at least 85,000 proteins per metagenome (Mediterranean Sea 30 m, Lake Baikal 20 m, Caspian Sea 15 m and Tous reservoir 12-25 m).

### Category classification of different proteins

Each protein was categorized in cytoplasmic/inner membrane, protein with transmembrane domains, proteins with signal peptides according to the SignalP [51] and Phobius [52] tools predictions. The pIs of the different proteins, their transmembrane domain topology and presence/absence of signal peptides for the eight microbes used in this comparison are shown in Supplementary Data Set S2.

### Structure homology-modelling and determination of the electrostatic surface potential of different proteins

The selected proteins for the pair microbe comparison were first modelled for their tertiary structure with the SWISS-MODEL online tool [53–55]. The extracted PDB was then visualized with PYMOL [56], and the electrostatic surface potential was calculated with APBS tool [57]. The surface potentials were mapped from −3 kcal mol^−1^ (red) to +3 kcal mol^−1^ (blue).

### Pan-genome analysis

The different freshwater and marine genomes used in the structural comparison were also subjected to a pan-genome analysis. Core and flexible genomes were determined with OrthoMCL and getHomologues softwares [58, 59].

## Supporting information

Supplementary Data Set S1

Supplementary Data Set S2

## List of abbreviations

MAG: Metagenome assembled genome
pI: Isoelectric point
ANI: Average Nucleotide Identity
AAI: Average Amino acid Identity
POCP: Percentage of conserved Proteins

## Authors’ contributions

FRV and PJCY conceived the project. PJCY performed the different analyses. FRV and PJCY wrote the manuscript.

### Acknowledgements

This work was supported by grant ‘VIREVO’ CGL2016-76273-P [AEI/FEDER, EU], (co-founded with FEDER funds).

