## Supplementary information, Figures S1-S10 for "Marine-freshwater prokaryotic transitions require extensive changes in the proteome"

[1] Evolutionary Genomics Group, Departamento de Producción Vegetal y Microbiología, Universidad Miguel Hernández, San Juan de Alicante, 03550 Alicante, Spain.

**Keywords:** Isoelectric point, marine-freshwater, proteome, salinity, electrostatic surface potential, charged amino acids

**Running title:** Marine-freshwater transitions proteome

**This PDF file includes:** Figs. S1 to S10.

**Other supplementary materials for this manuscript include the following:** Supplementary Data Sets S1-S2

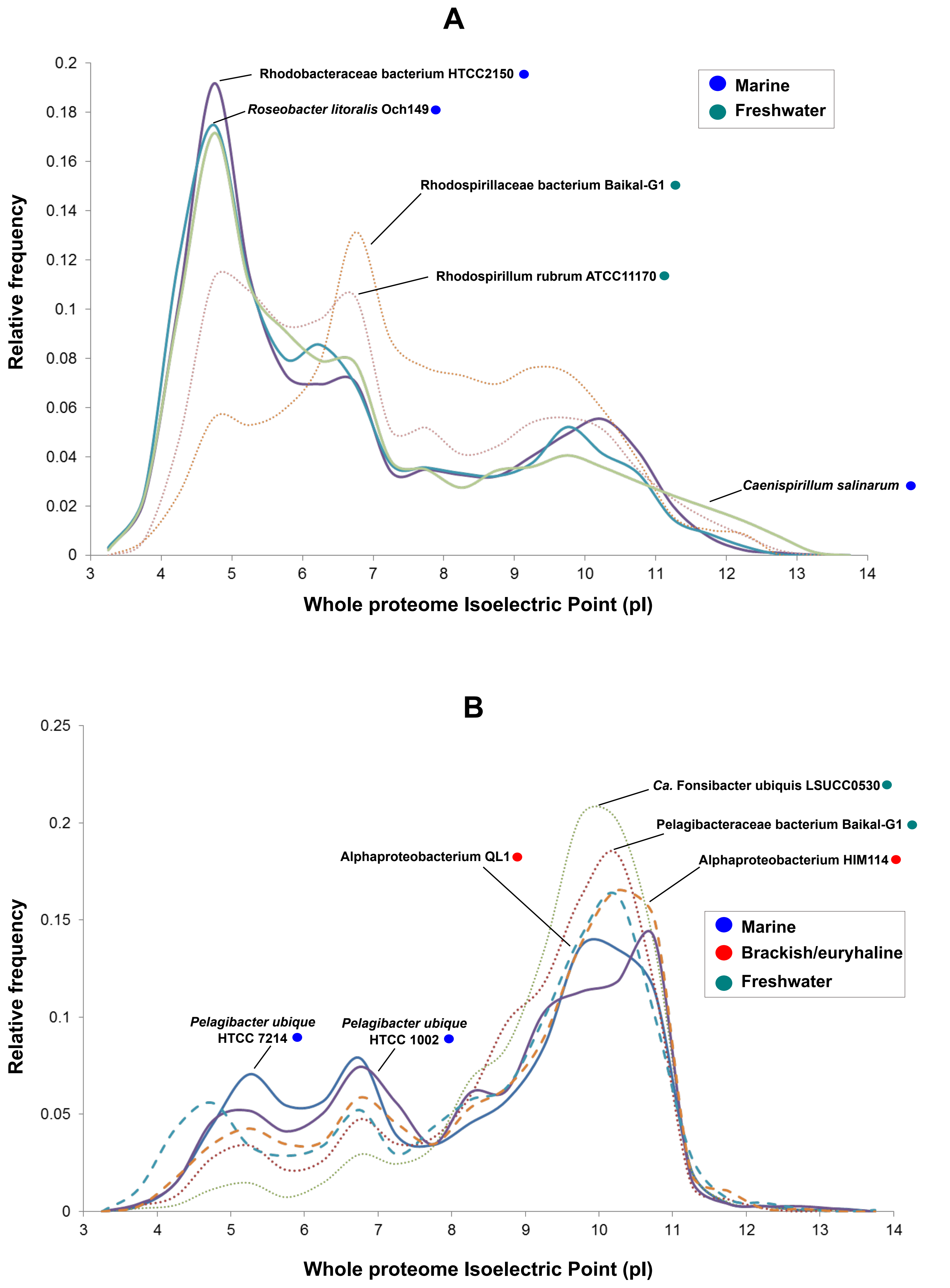


**Figure S1.** Whole proteome isoelectric point (pI) versus relative frequency of some representatives from different habitats of the class Alphaproteobacteria. A) Rhodospirillaceae and *Roseobacter* clades and B) SAR11 clades.

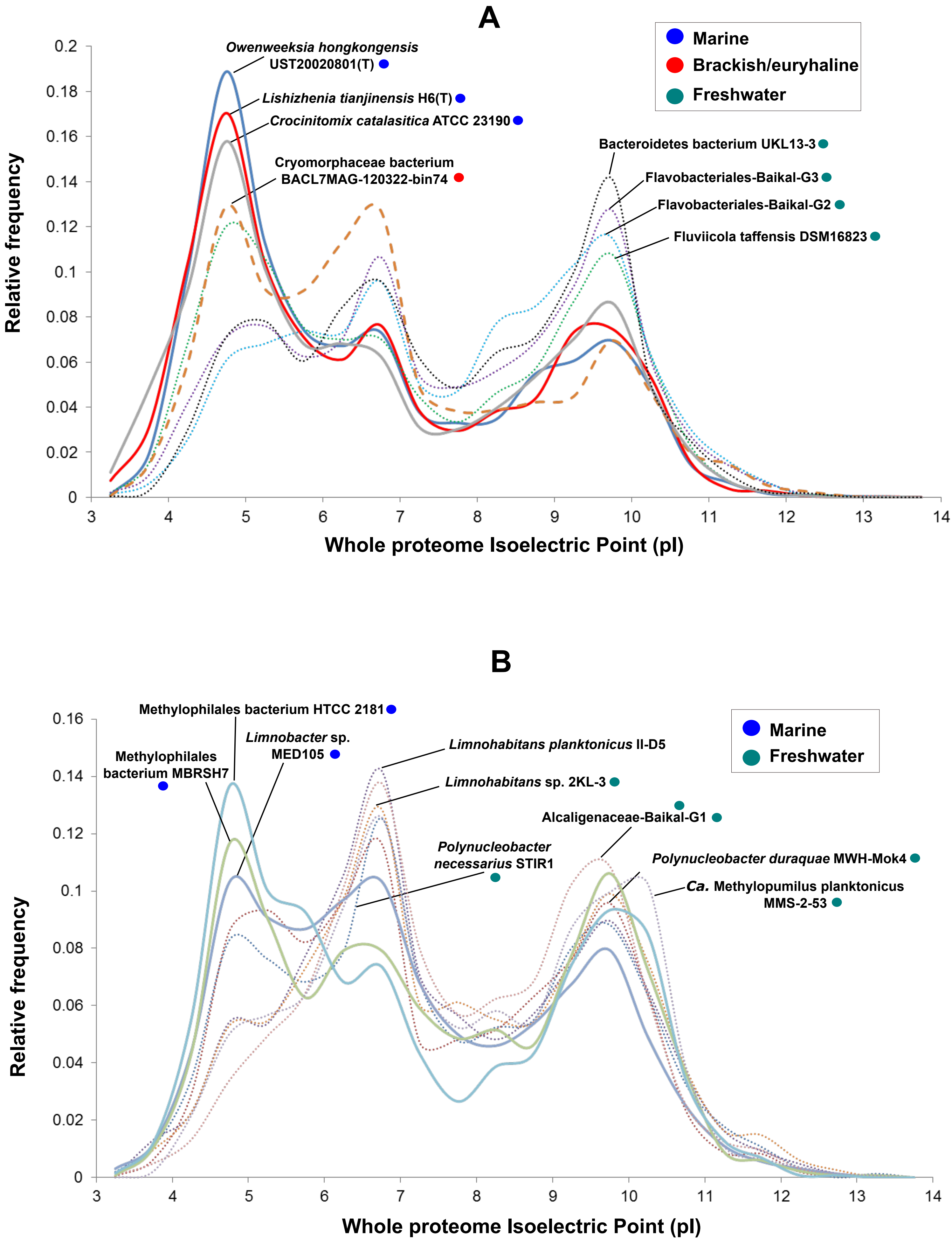


**Figure S2.** Whole proteome isoelectric point (pI) versus relative frequency of some representatives from different habitats of A) Bacteroidetes phylum and B) Order Betaproteobacterales.

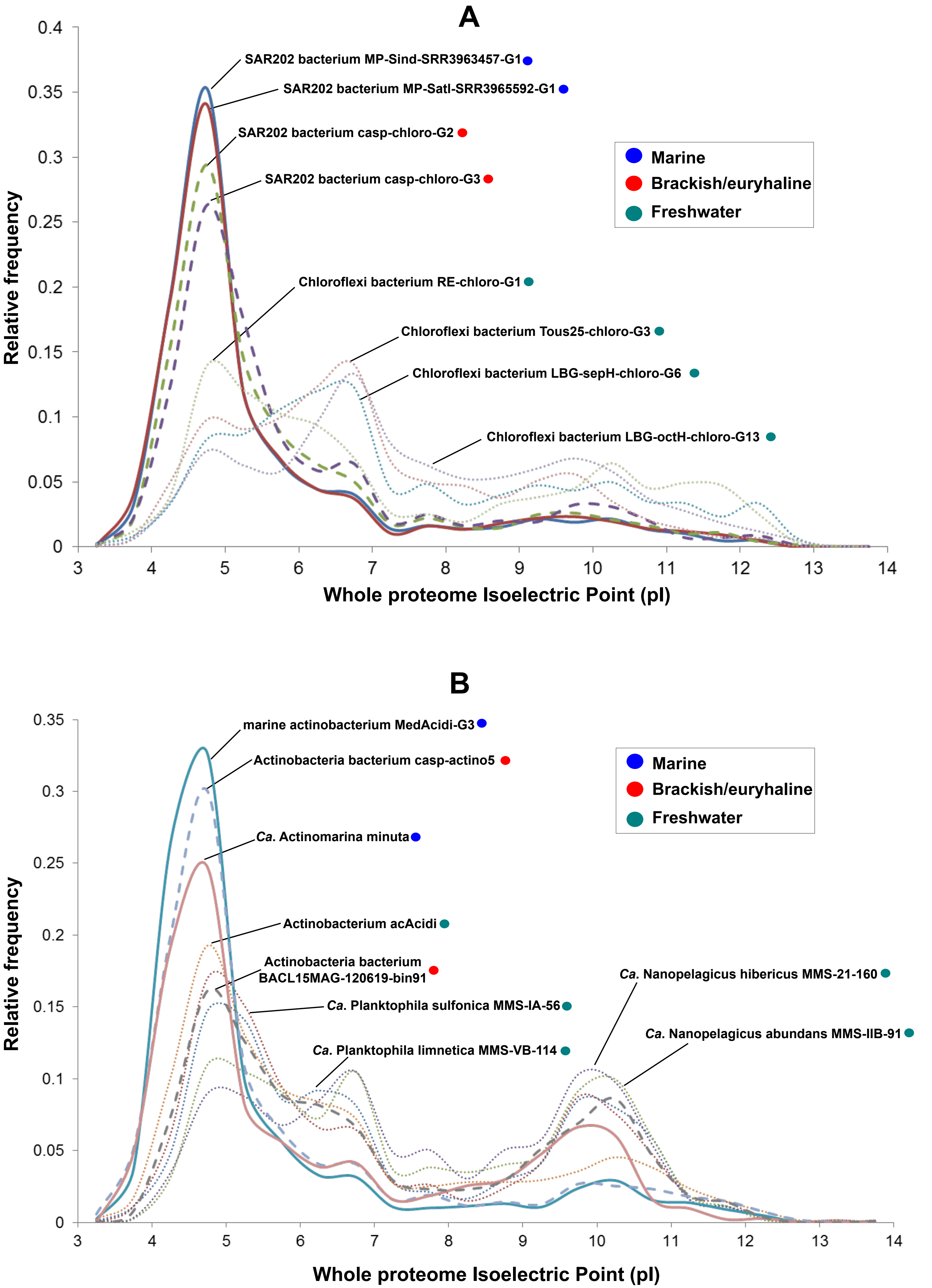


**Figure S3.** Whole proteome isoelectric point (pI) versus relative frequency of some representatives from different habitats of A) Chloroflexi phylum and B) Actinobacteria phylum.

## A

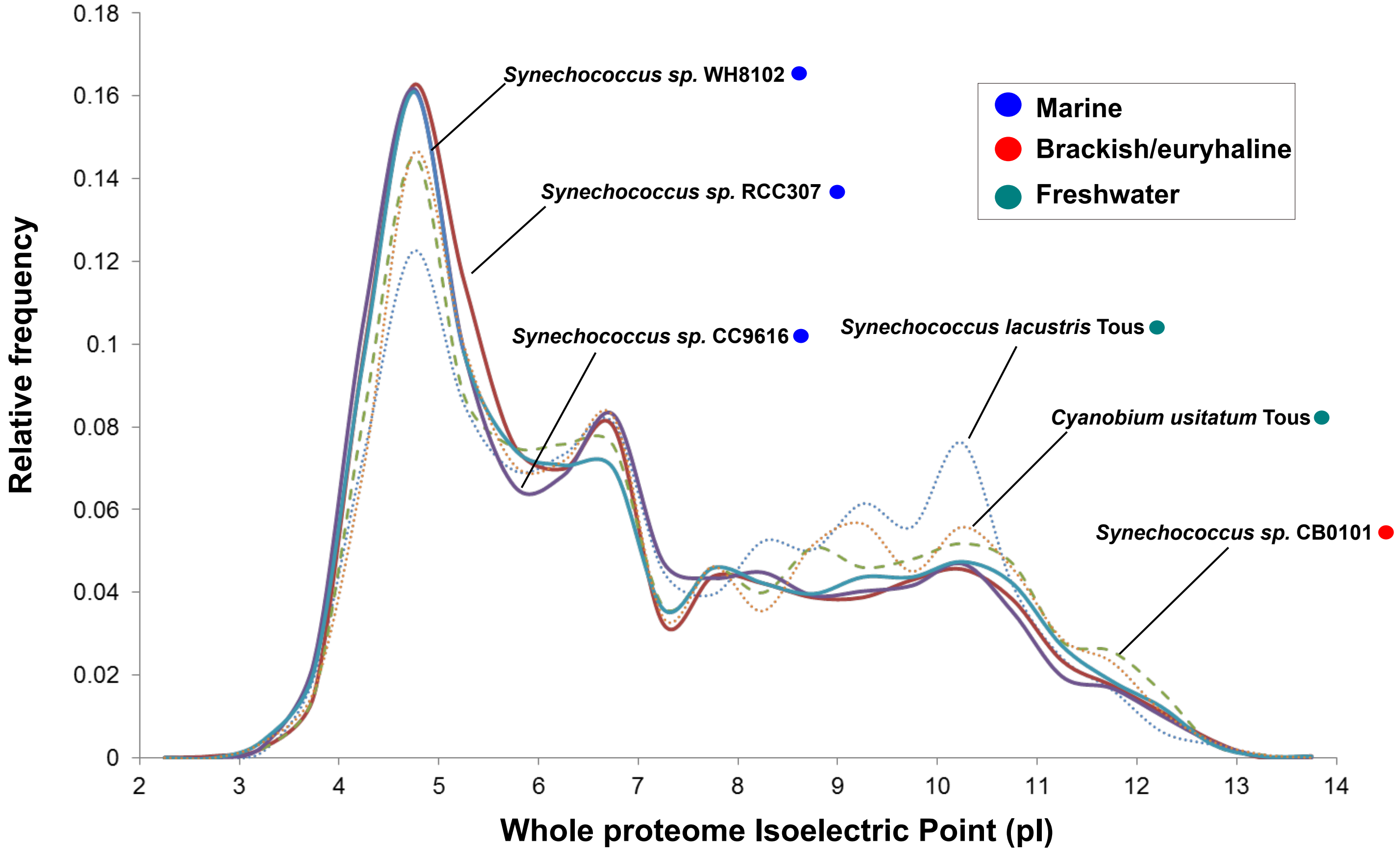


## B

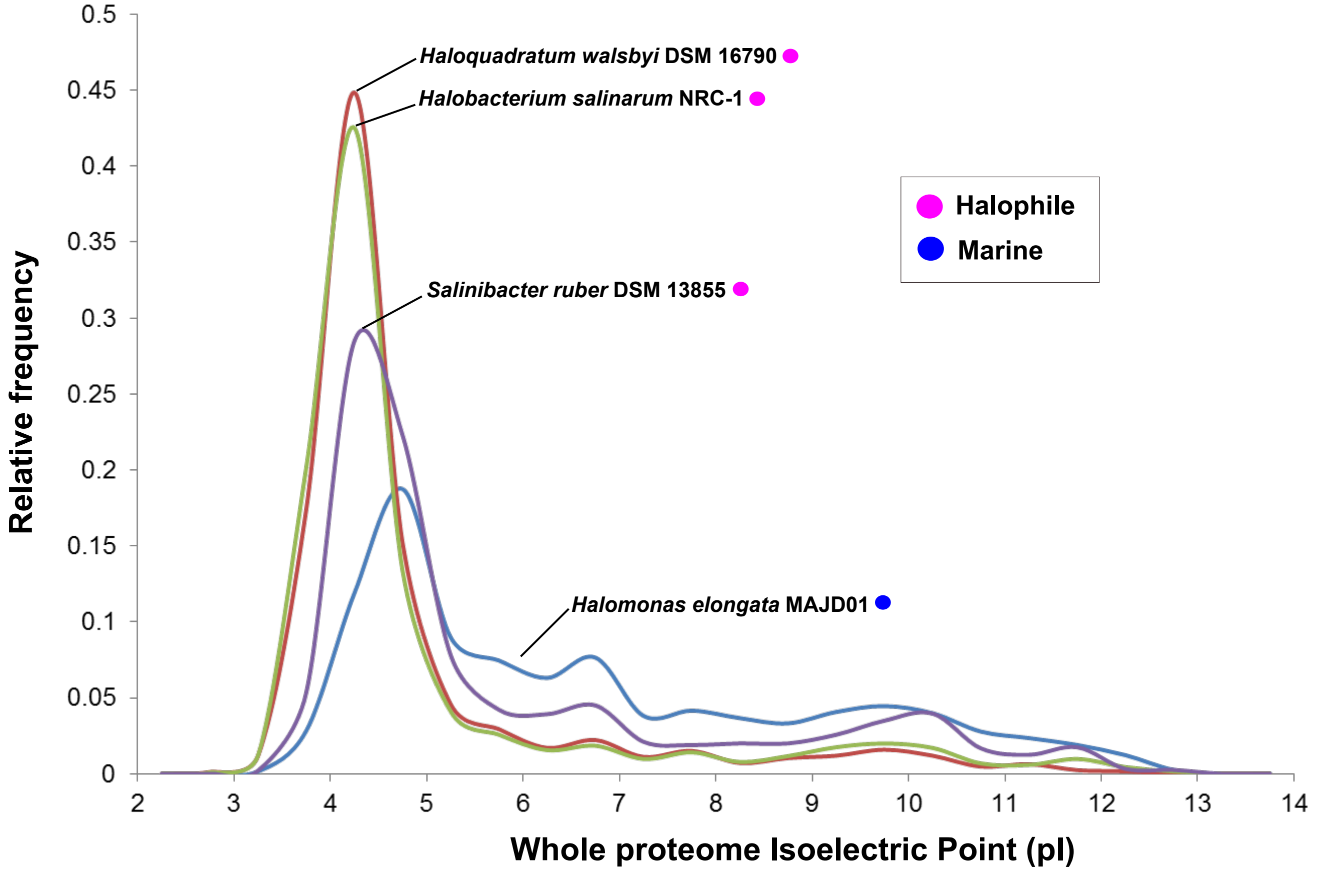


**Figure S4.** Whole proteome isoelectric point (pI) versus relative frequency of some representatives from different habitats of A) Genera *Synechococcus/Cyanobium* and B) Assorted halophiles (bacteria and acrchaea).

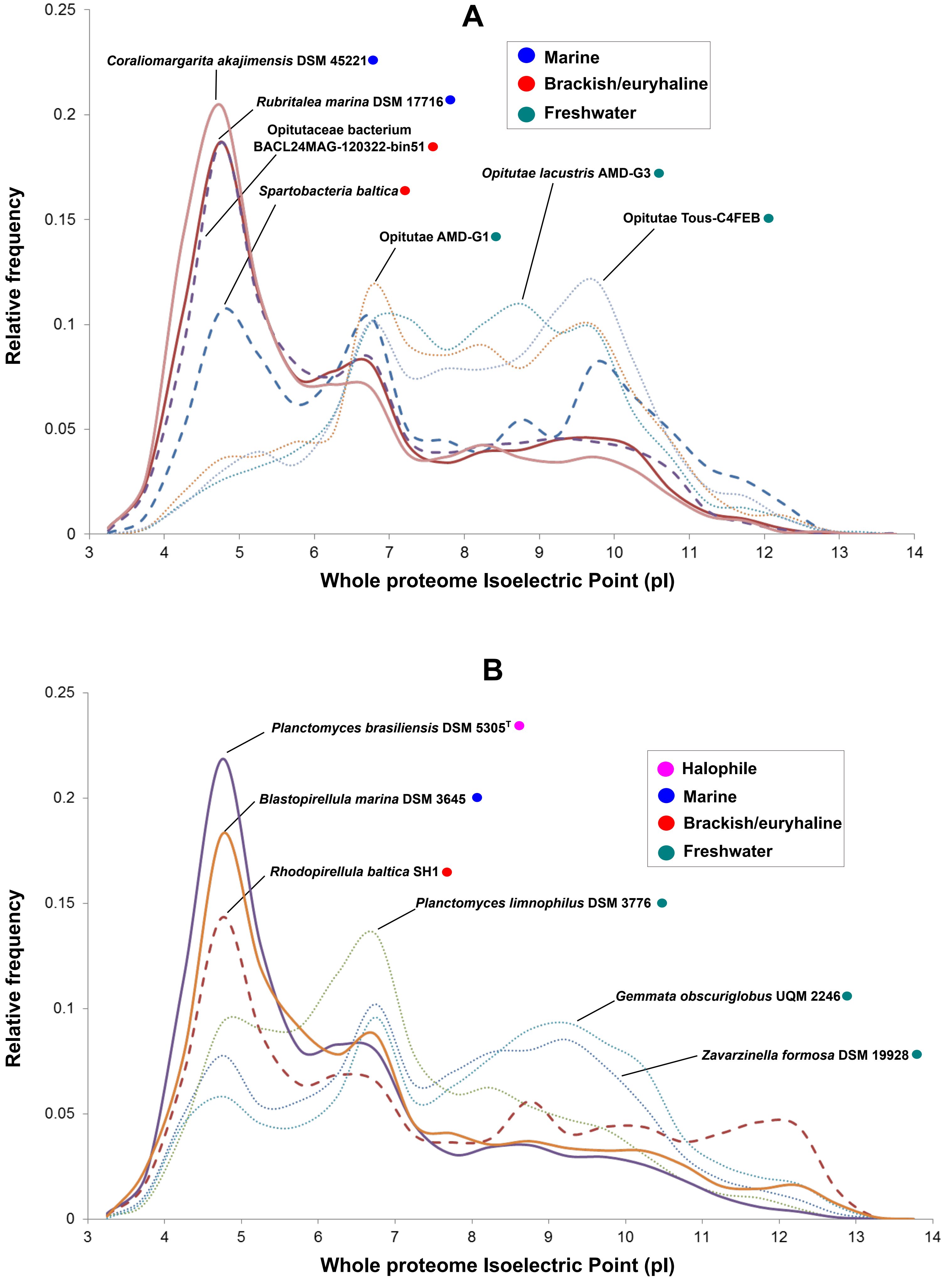


**Figure S5.** Whole proteome isoelectric point (pI) versus relative frequency of some representatives from different habitats of A) Verrucomicrobia phylum and B) Planctomycetes phylum.

## A

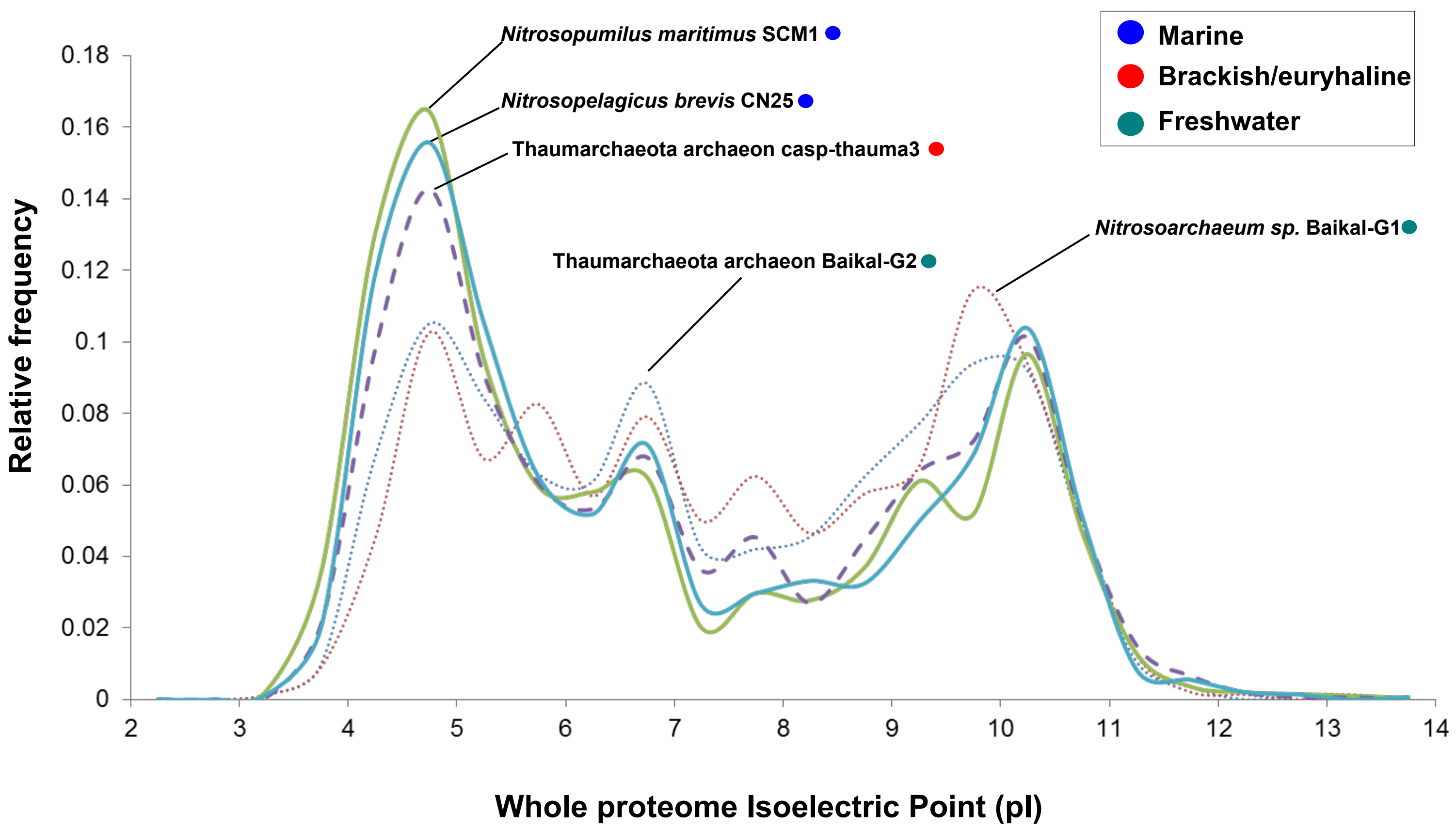


**Figure S6.** Whole proteome isoelectric point (pI) versus relative frequency of some representatives from different habitats of Thaumarchaeota phylum.

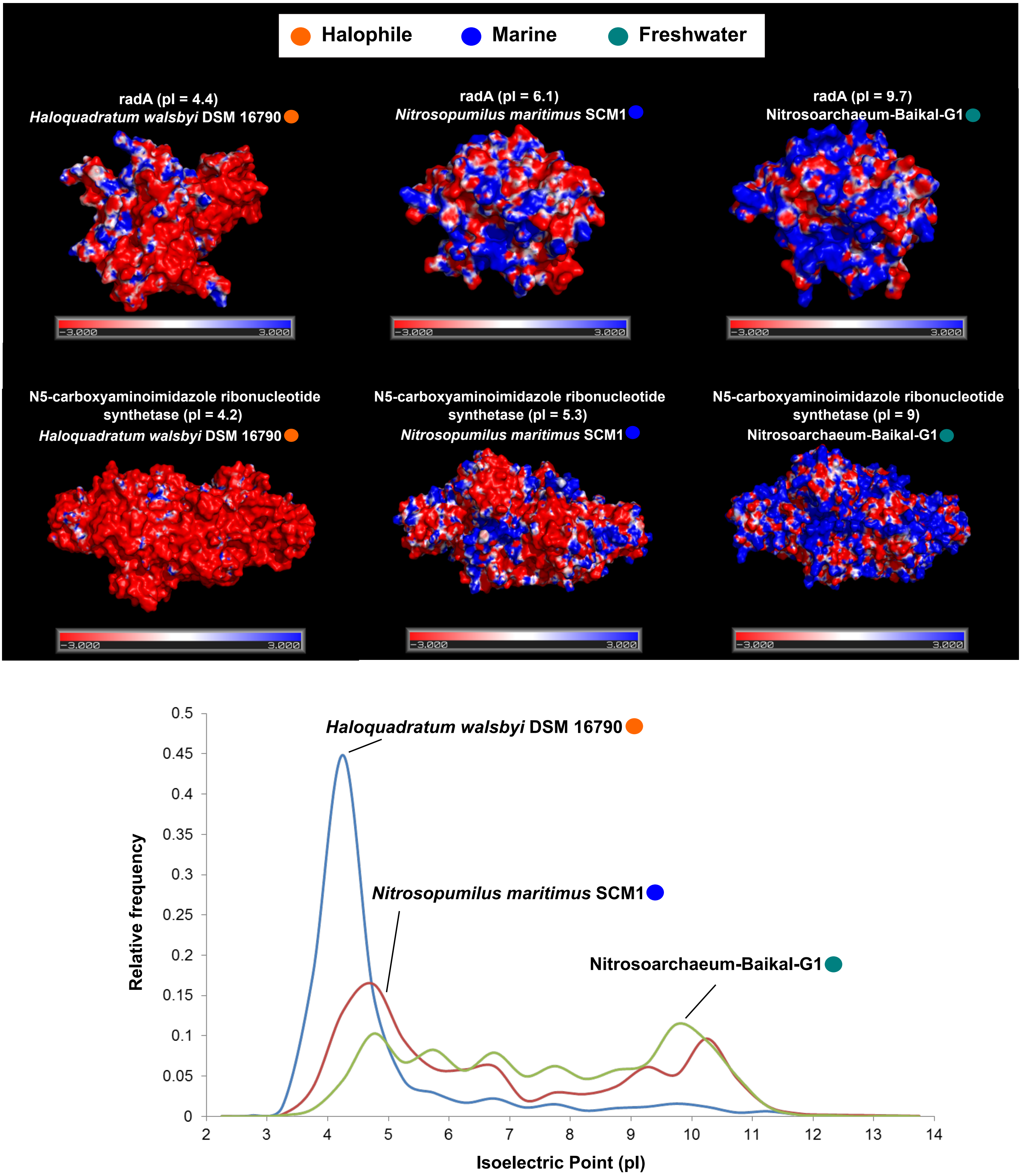


**Figure S7.** Structural model of proteins from different habitat-adapted Archaea. Tridimensional electrostatic surface potential plots of individual proteins selected for each archaeon. The potentials were mapped from -3 kcal mol-1 (red) to +3 kcal mol-1 (blue). Whole-proteome pI versus relative frequency of *Haloquadratum walsbyi* DSM 16790 (halophile), *Nitrosopumilus maritimus* SCM1 (marine), *Nitrosoarchaeum* sp. Baikal-G1 (freshwater).

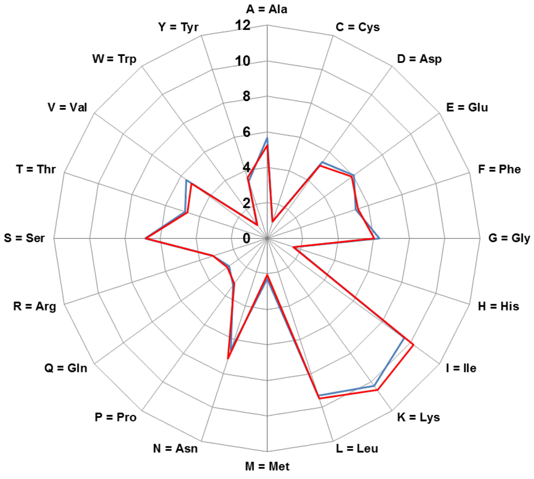


***Pelagibacteraceae* Baikal-G1**

***Pelagibacter ubique* HTCC 7214**

| Property | Residues | P.ubique HTCC 7214 (Mole %) | Pelagibacteraceae Baikal-G1 (Mole %) |
|---|---|---|---|
| Tiny | (A+C+G+S+T) | 24.689 | 23.791 |
| Small | (A+B+C+D+G+N+P+S+T+V) | 45.48 | 44.387 |
| Aliphatic | (A+I+L+V) | 30.134 | 30.152 |
| Aromatic | (F+H+W+Y) | 11.187 | 11.433 |
| Non-polar | (A+C+F+G+I+L+M+P+V+W+Y) | 52.627 | 52.274 |
| Polar | (D+E+H+K+N+Q+R+S+T+Z) | 47.373 | 47.726 |
| Charged | (B+D+E+H+K+R+Z) | 26.407 | 26.294 |
| Basic | (H+K+R) | 15.069 | 15.327 |
| Acidic | (B+D+E+Z) | 11.338 | 10.967 |

| Amino acid | *P.ubique* HTCC 7214 (Mole %) | Pelagibacteraceae Baikal-G1 (Mole %) |
|---|---|---|
| A = Ala | 5.652 | 5.221 |
| C = Cys | 1.001 | 0.989 |
| D = Asp | 5.302 | 5.07 |
| E = Glu | 6.036 | 5.897 |
| F = Phe | 5.252 | 5.398 |
| G = Gly | 6.33 | 6.035 |
| H = His | 1.593 | 1.555 |
| I = Ile | 9.552 | 10.191 |
| K = Lys | 10.28 | 10.565 |
| L = Leu | 9.317 | 9.472 |
| M = Met | 2.291 | 2.069 |
| N = Asn | 6.599 | 7.107 |
| P = Pro | 3.277 | 3.151 |
| Q = Gln | 2.66 | 2.78 |
| R = Arg | 3.196 | 3.208 |
| S = Ser | 6.869 | 6.817 |
| T = Thr | 4.838 | 4.728 |
| V = Val | 5.613 | 5.268 |
| W = Trp | 0.968 | 0.905 |
| Y = Tyr | 3.374 | 3.575 |

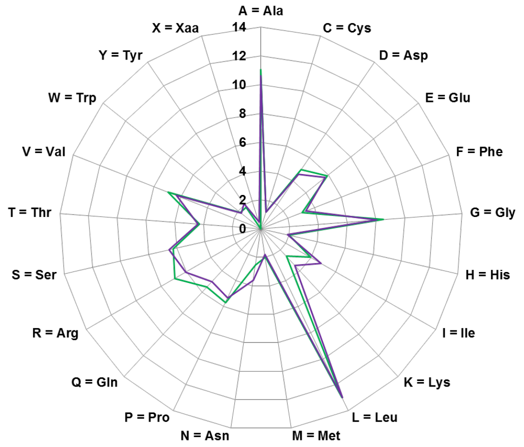


***Synechococcus lacustris* Tous**

***Synechococcus* sp. RCC307**

| Property | Residues | *Synechococcus* sp. RCC 307 (Mole %) | *Synechococcus lacustris* Tous (Mole %) |
|---|---|---|---|
| Tiny | (A+C+G+S+T) | 31.323 | 30.845 |
| Small | (A+B+C+D+G+N+P+S+T+V) | 51.438 | 50.679 |
| Aliphatic | (A+I+L+V) | 34.943 | 34.77 |
| Aromatic | (F+H+W+Y) | 8.7 | 9.042 |
| Non-polar | (A+C+F+G+I+L+M+P+V+W+Y) | 59.039 | 58.322 |
| Polar | (D+E+H+K+N+Q+R+S+T+Z) | 40.961 | 41.204 |
| Charged | (B+D+E+H+K+R+Z) | 22.394 | 21.638 |
| Basic | (H+K+R) | 11.53 | 11.363 |
| Acidic | (B+D+E+Z) | 10.865 | 10.275 |

| Amino acid | *Synechococcus* sp. RCC 307 | *Synechococcus lacustris* Tous |
|---|---|---|
| A = Ala | 11.07 | 10.624 |
| C = Cys | 1.219 | 1.222 |
| D = Asp | 4.959 | 4.592 |
| E = Glu | 5.906 | 5.683 |
| F = Phe | 3.057 | 3.325 |
| G = Gly | 8.509 | 8.043 |
| H = His | 2.016 | 1.873 |
| I = Ile | 3.953 | 4.796 |
| K = Lys | 2.61 | 3.463 |
| L = Leu | 12.985 | 13.017 |
| M = Met | 1.992 | 1.802 |
| N = Asn | 2.527 | 3.607 |
| P = Pro | 5.692 | 5.316 |
| Q = Gln | 5.513 | 5.003 |
| R = Arg | 6.904 | 6.027 |
| S = Ser | 6.244 | 6.546 |
| T = Thr | 4.282 | 4.41 |
| V = Val | 6.936 | 6.319 |
| W = Trp | 1.814 | 1.749 |
| Y = Tyr | 1.813 | 2.094 |
| X = Xaa | 0 | 0.474 |

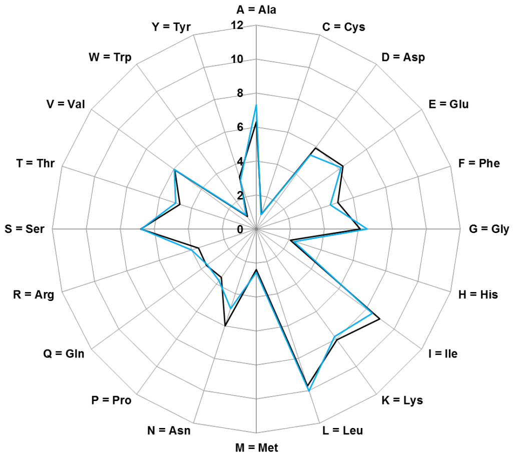


***Methylopumilus planktonicus* MMS-2-53**

**Methylophilales bacterium MBRSH7**

| Property | Residues | Methylophilales bacterium MBRSH7 (Mole%) | *Methylopumilus planktonicus* MMS-2-53 (Mole%) |
|---|---|---|---|
| Tiny | (A+C+G+S+T) | 24.848 | 26.443 |
| Small | (A+B+C+D+G+N+P+S+T+V) | 46.146 | 46.476 |
| Aliphatic | (A+I+L+V) | 30.884 | 31.73 |
| Aromatic | (F+H+W+Y) | 11.257 | 10.982 |
| Non-polar | (A+C+F+G+I+L+M+P+V+W+Y) | 52.989 | 54.054 |
| Polar | (D+E+H+K+N+Q+R+S+T+Z) | 47.011 | 45.946 |
| Charged | (B+D+E+H+K+R+Z) | 25.896 | 25.722 |
| Basic | (H+K+R) | 13.716 | 14.161 |
| Acidic | (B+D+E+Z) | 12.18 | 11.561 |

| Amino acid | Methylophilales bacterium MBRSH7 | *Methylopumilus planktonicus* MMS-2-53 |
|---|---|---|
| A = Ala | 6.294 | 7.302 |
| C = Cys | 0.92 | 0.887 |
| D = Asp | 5.891 | 5.396 |
| E = Glu | 6.289 | 6.166 |
| F = Phe | 5.047 | 4.587 |
| G = Gly | 6.117 | 6.506 |
| H = His | 2.085 | 2.341 |
| I = Ile | 8.965 | 8.437 |
| K = Lys | 8.045 | 7.824 |
| L = Leu | 9.695 | 10.021 |
| M = Met | 2.393 | 2.538 |
| N = Asn | 5.974 | 4.915 |
| P = Pro | 3.503 | 3.752 |
| Q = Gln | 3.624 | 3.561 |
| R = Arg | 3.587 | 3.996 |
| S = Ser | 6.805 | 6.758 |
| T = Thr | 4.713 | 4.99 |
| V = Val | 5.929 | 5.97 |
| W = Trp | 0.91 | 1.024 |
| Y = Tyr | 3.215 | 3.029 |

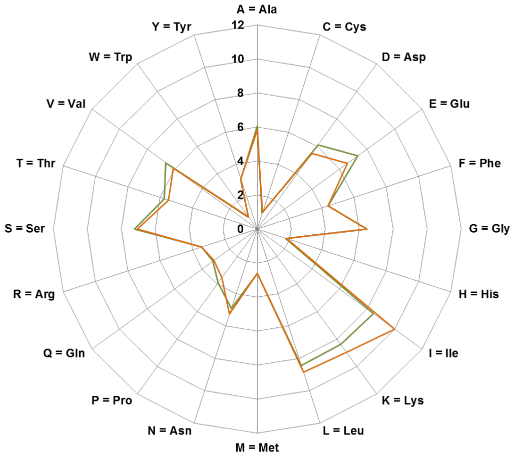


***Nitrosoarchaeum* sp. Baikal-G1**

***Nitrosopumilus maritimus* SCM1**

| Property | Residues | *Nitrosopumilus maritimus* SCM1 | *Nitrosoarchaeum* sp. Baikal-G1 |
|---|---|---|---|
| Tiny | (A+C+G+S+T) | 26.42 | 25.853 |
| Small | (A+B+C+D+G+N+P+S+T+V) | 47.982 | 46.256 |
| Aliphatic | (A+I+L+V) | 29.586 | 30.748 |
| Aromatic | (F+H+W+Y) | 10.147 | 10.168 |
| Non-polar | (A+C+F+G+I+L+M+P+V+W+Y) | 51.899 | 52.774 |
| Polar | (D+E+H+K+N+Q+R+S+T+Z) | 48.101 | 47.218 |
| Charged | (B+D+E+H+K+R+Z) | 27.047 | 26.295 |
| Basic | (H+K+R) | 13.586 | 14.204 |
| Acidic | (B+D+E+Z) | 13.461 | 12.091 |

| Amino acid | *Nitrosopumilus maritimus* SCM1 | *Nitrosoarchaeum* sp. Baikal-G1 |
|---|---|---|
| A = Ala | 6.055 | 5.792 |
| C = Cys | 0.983 | 1.056 |
| D = Asp | 6.124 | 5.509 |
| E = Glu | 7.337 | 6.582 |
| F = Phe | 4.403 | 4.382 |
| G = Gly | 6.445 | 6.47 |
| H = His | 1.772 | 1.824 |
| I = Ile | 8.461 | 10.006 |
| K = Lys | 8.393 | 8.952 |
| L = Leu | 8.432 | 8.855 |
| M = Met | 2.606 | 2.611 |
| N = Asn | 4.896 | 5.255 |
| P = Pro | 3.904 | 3.544 |
| Q = Gln | 3.222 | 3.133 |
| R = Arg | 3.42 | 3.428 |
| S = Ser | 7.199 | 7.05 |
| T = Thr | 5.738 | 5.485 |
| V = Val | 6.638 | 6.095 |
| W = Trp | 0.894 | 0.855 |
| Y = Tyr | 3.078 | 3.108 |

**Figure S8.** Star diagrams and amino acid composition of prokaryotic relatives from marine and freswhater origin.

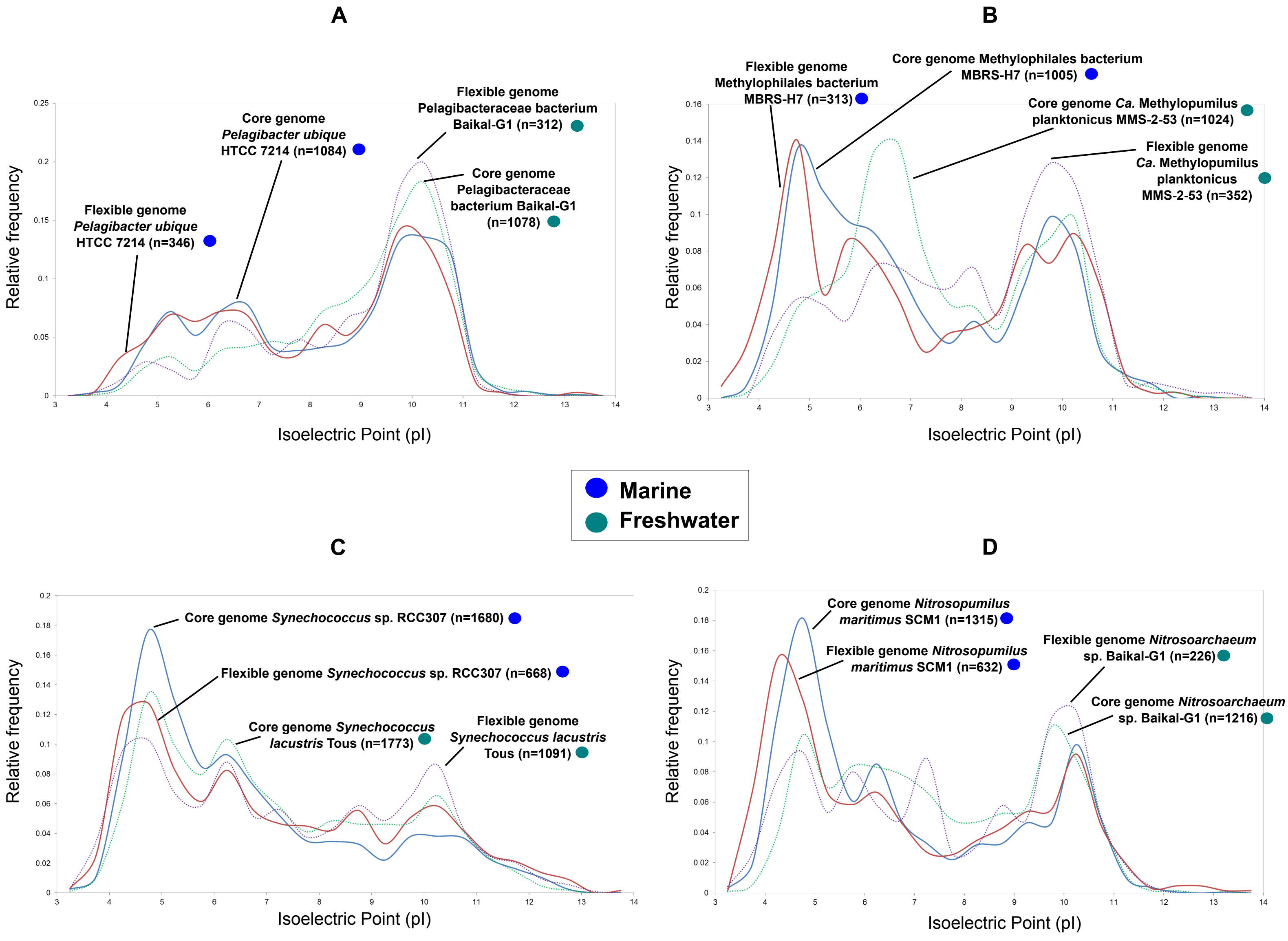


**Figure S9.** Isoelectric point (pI) versus relative frequency of the pan-genome (core and flexible genome) of freshwater and marine prokaryotes. N indicates the number of proteins accounting for either core or flexible genomes. A) *P.ubique* HTCC 7214 and Pelagibacteraceae bacterium Baikal-G1. B) *Ca*. Methylopumilus planktonicus MMS-2-53 and Methylophilales bacterium MBRS-H7. C) *Synechococcus* sp. RCC307 and *Synechococcus lacustris* Tous. D) *Nitrosopumilus maritimus* SCM1 and *Nitrosoarchaeum* sp. Baikal-G1.

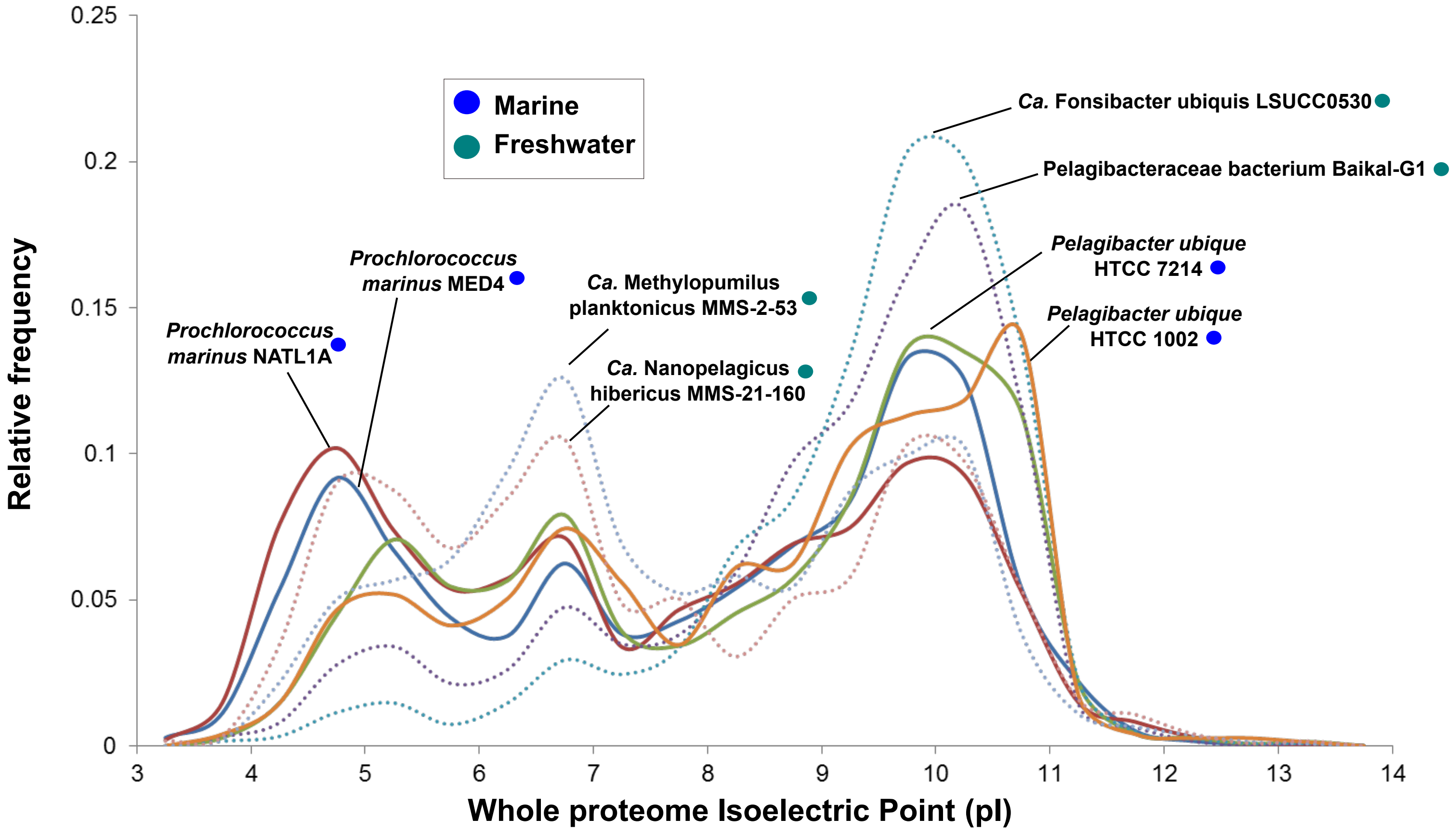


**Figure S10.** Whole proteome isoelectric point (pI) versus relative frequency of some streamlined bacteria from different habitats
